# Characterization of selective pressures acting on protein sites with Deep Learning

**DOI:** 10.1101/2025.07.01.662578

**Authors:** Estelle Bergiron, Luca Nesterenko, Julien Barnier, Philippe Veber, Bastien Boussau

## Abstract

It is often useful, in the field of molecular evolution, to identify the selective pressures acting on a particular site of a protein to better understand its function. This is typically done with likelihood-based approaches applied to codon sequences in a phylogenetic context. However, these approaches are computationally costly. Here we adapt a linear transformer neural network architecture, which has been shown to be able to reconstruct accurate pairwise distances from sequence alignments, to identify selective pressures acting on individual amino acid sites, in the mutationselection framework. We design different versions of the architecture and train and test them on simulations. We compare the results of one of our models to state-of-the-art likelihood-based methods and find that it outperforms them when it is applied to data that resemble its training data. Its performance decreases however when it is applied to datasets simulated with different parameter values. In all cases, our approach operates at a fraction of the computational cost of likelihood-based methods. These results suggest that such a neural network architecture can compare very favorably to state-of-the-art approaches to characterize selection pressures acting on coding sequences, but that it must be trained on datasets representative of empirical data.

## 1 Introduction

Deep learning approaches have been very successful in a range of scientific areas, and notably in genomics and biochemistry (Avsec et al., 2021; Jumper et al., 2021; Brandes et al., 2023). In recent years, they have begun to be applied in phylogenetics (Mo et al., 2024; Braichenko et al., 2025). First, they were used to reconstruct phylogenies with four taxa (e.g., Suvorov et al., 2019; Zou et al., 2020) but since then new models have been designed to work with arbitrary numbers of sequences (Nesterenko et al., 2025; Zhang et al., 2025; Blassel et al., 2026). In particular, the Phyloformer architecture (Nesterenko et al., 2025) was shown to be able to provide accurate estimates of the phylogenetic history of a simulated sequence alignment. Specifically, it performed very well with models of sequence evolution that include co-evolution, but its accuracy decreased as the number of sequences increased. Phyloformer is a deep neural network that takes in input an amino acid sequence alignment and infers a distance matrix, from which a phylogenetic tree can be obtained via existing methods. The architecture is based on several transformer blocks (Vaswani et al., 2017), implementing a custom linear attention mechanism that takes into consideration correlations between sites and between sequences to yield pairwise distance estimates.

We reasoned that such a network could provide estimates of selective regimes operating on protein sites, instead of estimates of distances between sequences. Inference of selective pressures acting on individual sites is usually based on the *d_N_ /d_S_* ratio between synonymous and non-synonymous codon substitution rates (Goldman and Yang, 1994; Muse and Gaut, 1994). When at a site *d_N_ /d_S_ <* 1, the ratio is indicative of negative or purifying selection; when *d_N_ /d_S_ ≈* 1, of neutral evolution; when *d_N_ /d_S_ >* 1, of positive selection. However, a different approach, that does not rely on *d_N_ /d_S_*, has also been proposed to model coding sequence evolution: mutation-selection models Halpern and Bruno, 1998; Yang and Nielsen, 2008. These models associate fitness parameters to the 61 non-stop codons, and thus enable detailed descriptions of the selective process, while separating it from the mutational process operating at the nucleotide level. This sophistication arguably results in a more realistic model of codon sequence evolution (Spielman and Wilke, 2015), but makes inference complicated (Rodrigue et al., 2010; Tamuri et al., 2012; Spielman and Wilke, 2016). However, Tamuri and dos Reis (2021) proposed a program for detecting sites that are under negative or positive selection along the entire phylogeny, in the mutation-selection framework. The program, swMutSel, operates in the penalized maximum likelihood framework, and remains unique in its ability. It is, however, costly to run. A deep learning approach for the same task may be significantly faster, but it is unclear how accurate it could be.

Previous works have demonstrated that deep learning could be used to predict characteristics of individual sites in an alignment. For example, Silvestro et al. (2024) have reported highly accurate substitution rates inferred from sequence alignments using a custom-designed recurrent neural network. Further, West et al. (2025) developed a convolutional neural network that can estimate whether a gene alignment with 8 sequences is under positive selection or not, when sequences have been simulated in the *d_N_ /d_S_* framework. Interestingly, they demonstrate that the network captures site-specific characteristics. These works suggest that deep learning approaches could provide useful alternatives to likelihood-based approaches for site-wise inference of selection.

In this paper, we report our efforts to build upon the Phyloformer architecture to classify protein sites as under neutral evolution or negative or positive selection. We train our network, SelRegAA, on simulations of protein alignments in the mutation-selection framework. In this framework, positive selection is simulated with the Persistent Positive Selection model of Tamuri and dos Reis (2021), which enforces that the fittest amino acid is always different from the current state. We investigate the performance of our network, its sensitivity to the parameters of the simulation, and its statistical calibration by modifying its structure and size. We compare its performance to swMutSel (Tamuri and dos Reis, 2021), the only available method to detect sites under positive selection that operates in the mutation-selection framework. However we also compare our approach to CODEML, a standard likelihood-based approach operating in the *d_N_ /d_S_* framework, therefore at a disadvantage on mutation-selection simulations. When simulations match the data used to train the network, we find that SelRegAA performs with an accuracy superior to CODEML, which, surprisingly, has itself a better accuracy than swMutSel. SelRegAA is orders of magnitude faster, and also faster than Godon which implements the same model as CODEML, but with a highly optimized code. However, the performance of SelRegAA decreases if it is applied to data that does not look like the data it has been trained on, in which case it obtains results similar to CODEML, or worse than it on simulations that match CODEML’s *d_N_ /d_S_* model. These results underline the importance of model and parameter choice when training a neural network on simulations. We conclude from these results that, in the future, a linear transformer network operating on codon sequences and trained on data representative of the range of alignments observed empirically, could perform very well, at a fraction of the cost of likelihood-based approaches.

## 2 Methods

The code for building, training, and testing the neural network is available here: https://gitlab.in2p3.fr/deelogeny/wp4/selregaa.

### 2.1 Neural network architecture

Our neural network is based on Phyloformer (Nesterenko et al., 2025), with two main differences (Fig. 1). First, we changed the output of the network. The original one outputs a distance matrix between sequences whereas in this setting we need site-wise annotations. Therefore, SelRegAA outputs a vector of 3 class probabilities per site of the alignment. The loss is computed as the cross-entropy between the output class probabilities and the true site-wise selection regime. Second, SelRegAA performs attention between sequences whereas Phyloformer performs attention between pairs of sequences.

**Figure 1:**
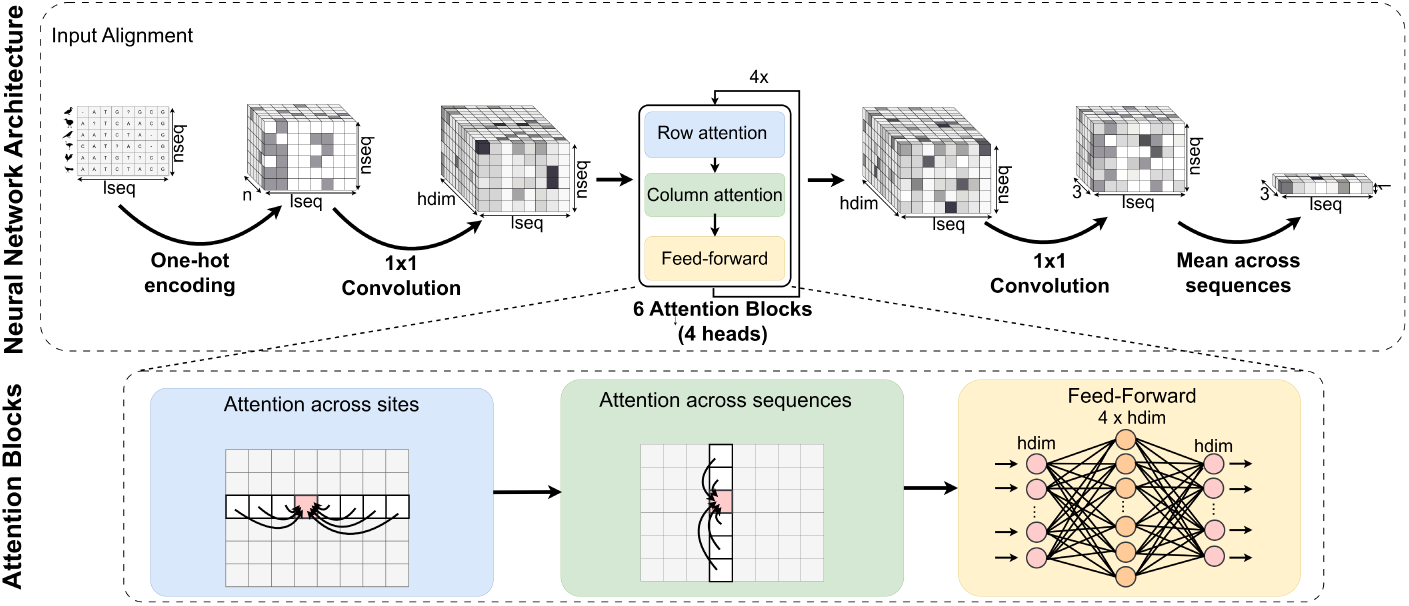
Structure of the base network. Top row: general architecture, from amino acid sequence encoding (left) to site-wise prediction of selective regimes (right). Bottom row: description of an attention block with 4 heads. lseq: number of sites in the alignment. nseq: number of sequences in the alignment. n: vector size after one-hot encoding (here, 21, for the 20 amino acids plus the gap character).

The input to the network is a one-hot encoding of an amino acid sequence alignment. Each amino acid in each sequence is thus transformed into a vector of size 21, corresponding to 20 amino acids, plus a gap character. These vectors are then converted into vectors of size 32, corresponding to the embedding dimension (see Supplementary Fig. 1). The value of these embedding vectors is updated as they traverse the network and its 6 attention blocks (Vaswani et al., 2017). At the end, the embedding vectors are converted into vectors of size 3, averaged per site and across sequences. These vectors of size 3 per site are used as predictors for neutral evolution, negative selection, and persistent positive selection.

### 2.2 Training algorithm

We trained our neural network with the AdamW optimizer with a learning rate of 0.001. We also used a scheduler with a 5 epochs warmup period followed by an exponential decay with a parameter *γ* of 0.99. The network was trained for a maximum of 200 epochs with an early stopping criterion, halting training if the validation loss value did not go down for 5 consecutive epochs. The network was trained with 10000 alignments, 90% of which for training, and 10% for validation.

### 2.3 Simulation protocol

#### 2.3.1 Tree simulation

Our simulation involves first simulating trees, and then simulating alignments along them. Trees were simulated as in Nesterenko et al. (2025). Briefly, ultrametric tree topologies were simulated using a Birth-Death process. Branch lengths were then altered to introduce heterogeneities between root-to-tip distances, and the trees were rescaled to match empirical tree diameters observed in the HOGENOM database (Penel et al., 2009) and the RaxML-Grove database (Höhler et al., 2021).

#### 2.3.2 Alignment simulation under a Mutation-Selection model

Codon alignments were simulated along the phylogenies using the sequence simulator Pastek (https://gitlab.in2p3.fr/pveber/pastek; tag=selregaa), using the multiselreg option. With this option, Pastek takes as arguments an effective population size, proportions of sites under negative or positive selection and neutral evolution, parameters of the nucleotide mutation matrix and of the insertion-deletion process, and alignment size. We use uniform echangeability and equilibrium frequencies for the nucleotide mutation matrix (*i.e.*, the Jukes-Cantor model (Jukes and Cantor, 1969)). For each alignment, an effective population size parameter is drawn from a uniform distribution between 0.5 and 5, and applied to all sites. For each alignment, the proportions of sites (in percents) under neutral evolution and under positive selection are both drawn from a Poisson distribution of rate 5 while the remaining sites are simulated under negative selection. We chose these values based on our expectation that most sites in empirical alignments are under purifying selection. For each alignment, gaps are simulated with a deletion rate drawn from 0.05, 0.1, 0.5 with weights 0.15, 0.7, 0.15, and according to the TKF91 model (Thorne et al., 1991). An insertion rate equal to 0 was chosen to avoid the difficulty of having to choose a selective regime for sites that appear further down along the tree. Pastek implements the Gillespie algorithm (Gillespie, 1977) to simulate sequences, and works as follows. For each site, Pastek randomly draws a selective regime category. All sites are simulated according to the Mutation-Selection model (Yang and Nielsen, 2008; Rodrigue et al., 2010), where synonymous codons are assumed to have the same fitness. Neutral sites are simulated by assuming a flat amino acid profile, where all amino acids have the same fitness. Negatively selected sites are simulated with an amino acid profile drawn from 263 empirically obtained profiles (Duchemin et al., 2023). Positively selected sites are simulated by drawing an amino acid profile as for negatively selected sites, but with an additional parameter *Z* to model persistent escape from the current amino acid (Tamuri and dos Reis, 2021). This parameter tends to increase the rate of substitutions away from the current state, irrespective of the current state. In our simulations, we used *Z* = 10, generating strong persistent positive selection; in their analysis of 3 gene alignments, Tamuri and dos Reis (2021) found that the *Z* values could reach values of more than 15, although rarely. To test for the sensitivity of the network to training parameters (see section 3.3.3), we also used another set of amino acid profiles, estimated from mammalian protein alignments (Latrille et al., 2024). From this set, we randomly picked two samples of 250000 profiles (profiles0 and profiles1) from 1000 alignments. Pastek can produce codon alignments and the corresponding amino acid alignments. The amino acid alignments were used to train and test the neural networks and to run swMutSel (Tamuri and dos Reis, 2021), and the codon alignments were used with CODEML (Yang, 2007) and Godon (Davydov et al., 2017). The resulting alignments are 250 sites long and contain 50 sequences.

#### 2.3.3 Alignment simulation under a *d_N_ /d_S_* model

To evaluate the sensitivity of SelRegAA to a strong mismatch between the evolutionary model used for training and that used to generate the test data, ten codon-sequence alignments were simulated using the *d_N_ /d_S_* model implemented in INDELible. The simulations used phylogenetic trees generated as above. Each alignment comprised 250 codon sites. Sites were assigned to three selective regimes, with 90% evolving under purifying selection, 5% under neutral evolution, and 5% under positive selection. For each alignment, we chose values for the *d_N_ /d_S_* ratio, *ω*, as in West et al. (2025). The *ω* assigned to sites under purifying selection was drawn from a uniform distribution between 0.1 and 0.9. Neutral sites were assigned a fixed value of *ω* = 1, whereas the value assigned to positively selected sites was drawn from a uniform distribution between 1.5 and 5. The transition-to-tranversion rate ratio, *κ*, was independently chosen for each alignment from a uniform distribution between 2 and 3. The resulting codon alignments were translated into amino acid alignments for analysis with SelRegAA, whereas the original codon alignments were used for analyses with CODEML.

### 2.4 Comparison between SelRegAA and likelihood-based methods

#### Alignments for method comparison

To compare results and computing time, we applied competing methods to 10 different alignments. Two sets of 10 alignments were produced, corresponding to two different amino acid profiles, the default 263 profiles from Duchemin et al. (2023) (default263) and one of our two sets of profiles estimated from mammalian alignments (profiles0).

#### Running swMutSel

The persistent positive selection test was performed using swMutSel as described by Tamuri and dos Reis (2021). First, mutation parameters (nucleotide frequencies and transition/transversion ratio) and branch lengths were estimated for each alignment using CODEML. Then, a null mutation-selection model and an alternative model with persistent-positive selection were fitted using swMutSel, with mutation parameters and branch lengths fixed. Site-specific amino acid fitness profiles were estimated with Dirichlet regularization (*α* = 0.01). The alternative model with persistent positive selection (PPS) was fitted at each site using swMutSel and had an additional exponential penalty applied to the PPS parameter (*λ* = 0.001). A likelihood ratio was then calculated for each site, between the null and alternative models. Statistical significance was assessed using parametric bootstraping (Cox test): sequence alignments were simulated under the null model using swMutSel, and the two models were refitted to each site to obtain an empirical null distribution of the likelihood ratio. P-values were computed from this distribution. Multiple testing correction was then performed separately for each alignment using the Benjamini-Hochberg procedure, and a significance threshold of 0.05 was applied.

Data, configuration and results of each swMutSel evaluation are available in the swmutsel folder of https://gitlab.in2p3.fr/deelogeny/wp4/selregaa.

#### Running CODEML

CODEML, from PaML version 4.10.6, was applied both under the M2 model (NSSites = 2) and under the M8 model (NSSites = 8). Site-wise prediction of selection regimes was made using the Bayes Empirical Bayes method (Yang et al., 2005). Data, configuration and results of each CODEML evaluation are available in the CODEML folder of https://gitlab. in2p3.fr/deelogeny/wp4/selregaa.

#### Running Godon

Godon was run under the M8 model to the same set of alignments. Sites under positive selection were identified using the Bayes Empirical Bayes approach. Configuration for Godon evaluations is available in the Godon folder of https://gitlab.in2p3.fr/deelogeny/wp4/selregaa.

### 2.5 Estimation of computing time

Networks were trained on one V100 GPU. To compare computing times, CODEML, Godon and our base model were run on a cluster of computers, containing Intel Haswell, Broadwell, Skylake and AMD Epyc processors, using one CPU per run. As a result, computations did not all run on the exact same architecture and cannot be compared accurately. However, the orders of magnitude differences in running times that we witnessed would likely remain if measured on a standardized configuration.

### 2.6 Performance metrics

To evaluate the performance of our networks, we report precisions, recalls and *F*_1_ scores. The precision for a class is the proportion of correctly predicted sites among all the sites predicted by the network to belong to that class. The recall for a class is the proportion of correctly predicted sites among all the sites in the dataset belonging to that class. The *F*_1_ score for a class is the harmonic mean of the precision and recall scores for that class. It ranges between 0 and 1.0, with 1.0 for a perfect prediction with a recall and a precision of 1.0. As a global performance metric we use a global *F*_1_ score as an average of the *F*_1_ scores of each class.

### 2.7 Analysis of empirical alignments

We evaluated SelRegAA on three empirical protein-coding sequence datasets previously analyzed by Tamuri and dos Reis (2021) : the hemagglutinin protein (HA) from human influenza H1N1 virus, the rbcL protein subunit from flowering plants (rbcL), and the mitochondrial cytochrome b (CYTB) protein from mammals.

Positively selected sites were identified using SelRegAA, CODEML model M8 with Bayes Empirical Bayes inference, and a proxy derived from the PPS model.

#### CODEML

CODEML analyses were performed using the same settings as in section 2.4. For the HA alignment specifically, cleandata = 1 was used to prevent an error caused by gaps or ambiguous character. With this setting, CODEML performs complete deletion of codon columns containing at least one gap or ambiguity character. CODEML site coordinates were mapped back to the coordinates of the original alignment before comparisons between methods.

#### *ω*-based proxy derived from the PPS model

To our knowledge, the complete results from the PPS test were not available from the original publication (Tamuri and dos Reis, 2021). We therefore approximated PPS test predictions using the estimated site-specific ratio of nonsynonymous to synonymous substitution rates (*ω_k_*), classifying sites with *ω_k_ >* 1 as positively selected. Estimates obtained with the weak regularization setting *λ* = 0.001 were retained. *λ* is the regularization parameter controlling the exponential penalty applied to *Z_k_*, the persistent postive selection component of the model. As shown in the analysis of Tamuri and dos Reis (2021), the proxy used here may identify fewer positively selected sites than the full PPS test because several sites detected by the PPS test had *ω_k_ <* 1.

## 3 Results

Detailed results for each experiment are available at https://deelogeny.pages. in2p3.fr/wp4/selregaa/. All models are tested on unseen test data.

### 3.1 Network architecture and training

In this section we compare the performances of our base model to variants of it, where the training or the architecture of the network were changed.

#### 3.1.1 Training parameters

Our base model was trained on 10000 alignments. We performed 4 independent trainings of our base network architecture to evaluate the reproducibility of our training. Supplementary Table 1 shows that all replicate trainings resulted in very similar performances, with *F*_1_ scores ranging between 0.759 (model base 2) and 0.769 (model base 1). Models 3 and 4 obtained *F*_1_ scores of 0.762 and 0.763. In the rest of the manuscript, we will use model base 1 as our default model, and will name it SelRegAA. Due to our stopping criterion, training times varied, with model base 1’s training taking 1h43 minutes for 69 epochs, model base 2’s training taking 54 minutes for 38 epochs, model base 3 1h10 for 45 epochs, and model base 4 1h47 for 49 epochs. This experiment suggests that differences of around 0.01 in *F*_1_ scores can be observed between replicates. In terms of precision and recall, the largest differences are observed for classifying sites under positive selection, reaching around 0.03.

We explored the network’s performance when it is trained on a larger number of sequences, i.e. 50000 alignments instead of 10000. After training on the larger dataset, the network obtains an *F*_1_ score of 0.788, better than the 0.769 of base model 1/SelRegAA.

We note that both precision and recall are improved for detecting positively selected sites with the larger training dataset size (Supp. Table 2). In the following tests, we train the variants of our network on 10000 alignments to avoid large computational footprints.

**Table 1:**
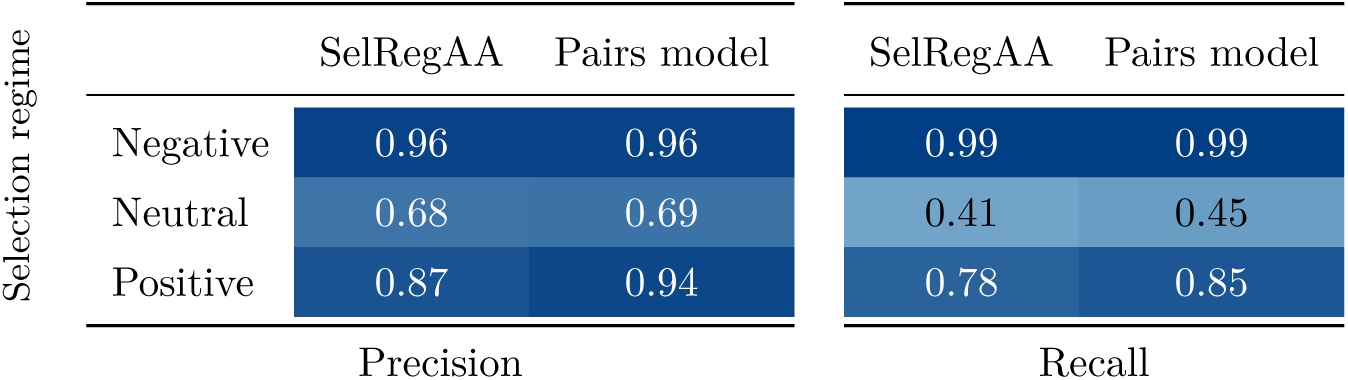
A sequence level model performs less well than a costly pair-level model.

**Table 2:**
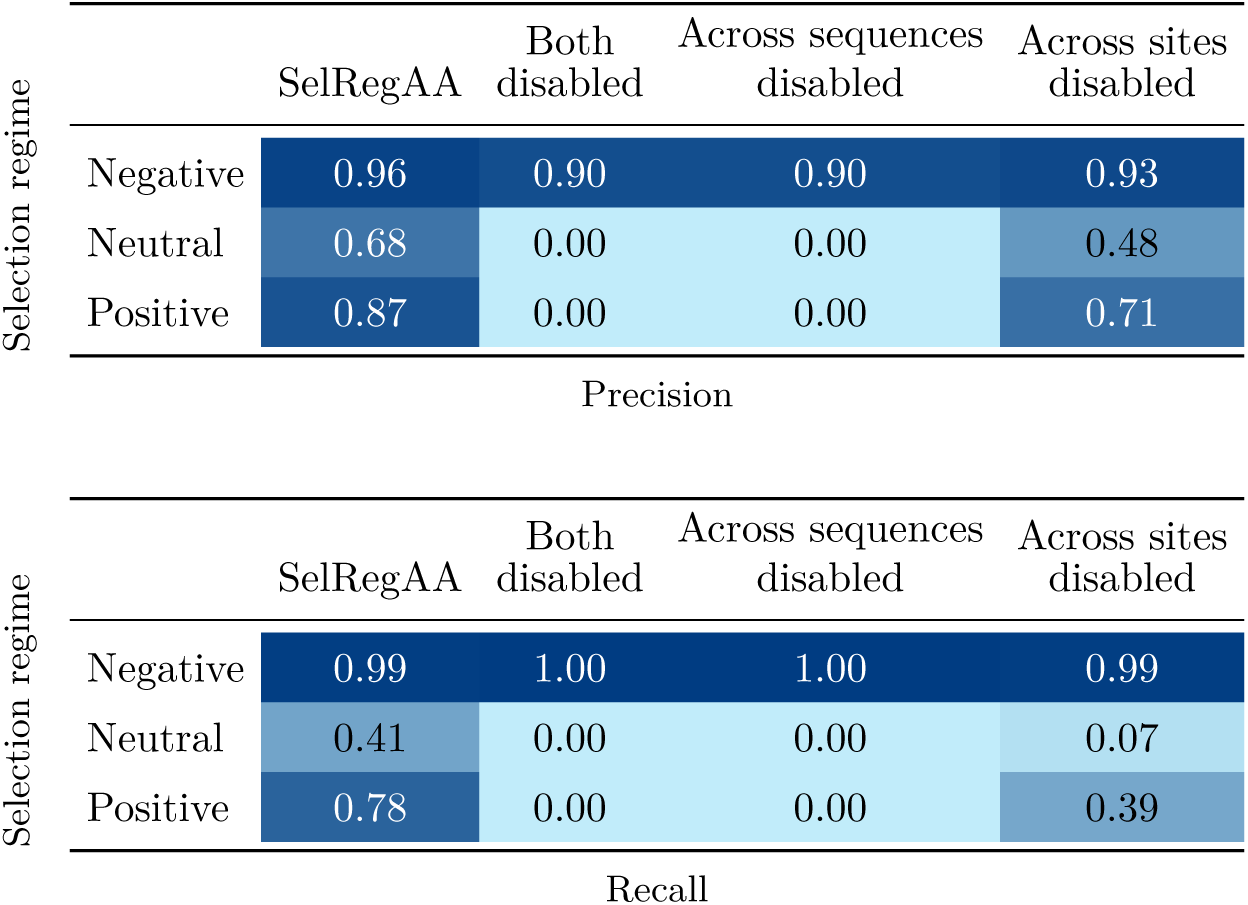
Effect of attention across sequences and across sites on precision and recall. Compared to SelRegAA (left), removing the two axial attention mechanisms degrades performance. When both attention mechanisms are disabled or when only attention across sites is used, the network fails to predict site categories (second and third columns). When only attention across sequences is used, the network’s performance is decreased markedly (fourth column).

#### 3.1.2 Varying the network architecture

Our base network uses attention between sequences and thus differs from Phyloformer, which works on pairs of sequences to provide accurate estimates of pairwise distances. Attempts at working at the sequence level to reconstruct pairwise distances had been much less accurate (Nesterenko et al., 2025). However, working on pairs of sequences entails a memory footprint that scales quadratically with the number of sequences in the alignment, compared to linearly when working with sequences. We investigated the performance of an architecture operating on pairs of sequences, and compared it to SelRegAA, which works on individual sequences.

Table 1 shows that the pair model performs better than the sequence model, in particular for the detection of positive selection, with a gain in precision and recall of 0.07. These gains appear meaningful since variation in these metrics between our four replicates (Sup. Table 1) were within a range of 0.03. They likely result from the gain in expressivity associated with considering pairs instead of individual sequences, and not from the number of parameters, because the sequence model has 102, 051 parameters, and the pair model has 68, 323 parameters. The pair model has fewer parameters because only 4 attention blocks were used instead of 6, to allow a batch size of 8 alignments during training. However, training the pair model was 15 times slower than the sequence model, and had a markedly larger memory footprint. In the following, we choose to focus on the sequence model because it has the best potential for scaling up to large datasets, where fast approaches are most needed.

##### Size of the embedding

The base network takes a sequence alignment as input, in which all sites are one-hot encoded into vectors of dimension 21. In turn, these vectors are embedded into vectors of size 32, that then pass through the attention blocks. We investigated the impact of changing the size of the embedding. Supplementary Table 3 shows that the size of the embedding has little influence on the performance of the network. Decreasing it to 16 instead of 32 results in a slightly lower *F*_1_ score of 0.749, and increasing it to 48 results in the same *F*_1_ score of 0.769.

##### Number of attention blocks and heads

We investigated a range of network sizes, by changing the number of attention blocks and attention heads in our network.

Our base model has 6 attention blocks that each combine column and row attention mechanisms. We tested reducing this number to 2 and 4, or increasing it to 8. Supplementary Table 4 shows that the performance of the network is not very sensitive, if at all, to the number of attention blocks.

Inside each axial attention mechanism, we use several heads whose roles are to focus on different characteristics of the data. Our base model has 4 attention heads. We trained alternative models with 2 or 6 attention heads, keeping head size constant (size 8). As a result, embeddings become larger with more heads: the network with 2 heads uses embeddings of size 16, and the network with 6 heads uses embeddings of size 48. Supplementary Table 5 shows that the number of heads affects the performance of the network slightly. We observe a very weak decrease in performance when only 2 heads are used (*F*_1_ score of 0.750 instead of 0.769 for base model 1/SelRegAA), but no improvement when 6 heads are used (*F*_1_ score of 0.768).

##### The effects of row and column attentions

The Phyloformer architecture performs attention across both sites and sequences in each attention block. To investigate the impact of each type of attention on site-wise classification accuracy, we trained two networks in which either attention had been deactivated and one network in which both attention mechanisms had been deactivated. Table 2 shows that both attention mechanisms are important. Attention across sites is important, as removing it degrades the *F*_1_ score from 0.769 for SelRegAA to 0.527. Just removing the attention across sequences is enough to completely destroy the network’s ability to predict site categories: the network then predicts that all sites belong to the “negative selection” category, given that it is the category with the largest number of sites in the data.

### 3.2 Sensitivity to dataset characteristics

#### 3.2.1 Sensitivity to alignment size

Our network was trained on datasets with 50 sequences and 250 sites. We investigated its performance on alignments with larger numbers of sites and sequences. When trained and tested on alignments with 100 sequences instead of 50, we observe that the network’s recall increases both for the neutral and positive selection categories, with a better precision for positive selection sites (Table 3).

**Table 3:**
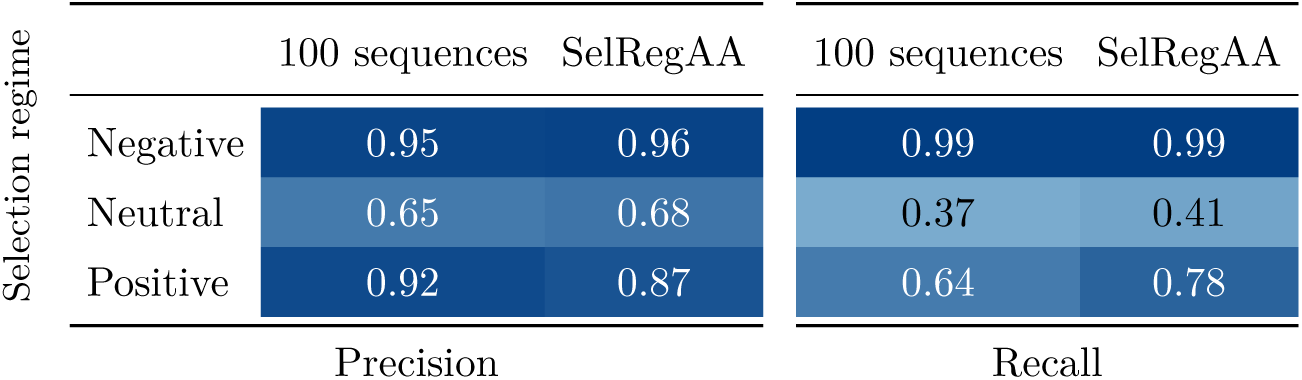
The trained network has better performance when it is trained and tested on alignments with a larger number of sequences.

When trained on alignments with 400 sites instead of 250, the performance of the network does not improve (Supp. Table 6). A larger number of sites does not result in an improved prediction accuracy per site. Increasing the number of sites in the alignment can only be useful for more accurately accounting for the phylogenetic structure in the data, which does not seem to matter here. Conversely, since the prediction is made at the level of individual sites, a larger number of sequences provides more information per site, resulting in a higher performance. Interestingly, the higher performance obtained on alignments with 100 sequences does not propagate to smaller alignments: the network trained on large alignments performs less well on alignments with 50 sequences than the network trained on smaller alignments (data not shown).

#### 3.2.2 Weighting the loss to account for imbalance in the proportions of the categories

The training data was generated with 5% of the sites under neutral evolution, 90% under negative selection, and 5% under persistent positive selection, on average per alignment. This strong imbalance in the proportions of the different categories could lead the network to focus on accurately predicting sites under negative selection, with a poorer performance for the other two minor categories. To investigate the strength of this effect, we trained a network with a weighted loss on the same training data. This weighted loss effectively penalizes errors in the minor categories more than errors in the negative selection category, by weighting errors with the inverse of each class frequency. The weights were 0.378 for negative selection and 6.66 for the other two categories. Table 4 shows that the model trained with weighted loss has better recall in the minor categories and a higher precision in the most frequent category. However, the precision in the minor categories is markedly decreased. Overall, the performance of the weighted model is reduced, with an *F*_1_ score of 0.666 compared to 0.769 for SelRegAA. This is not unexpected as the *F*_1_ score depends on class balance, and in the following we keep our focus on the unweighted model SelRegAA.

**Table 4:**
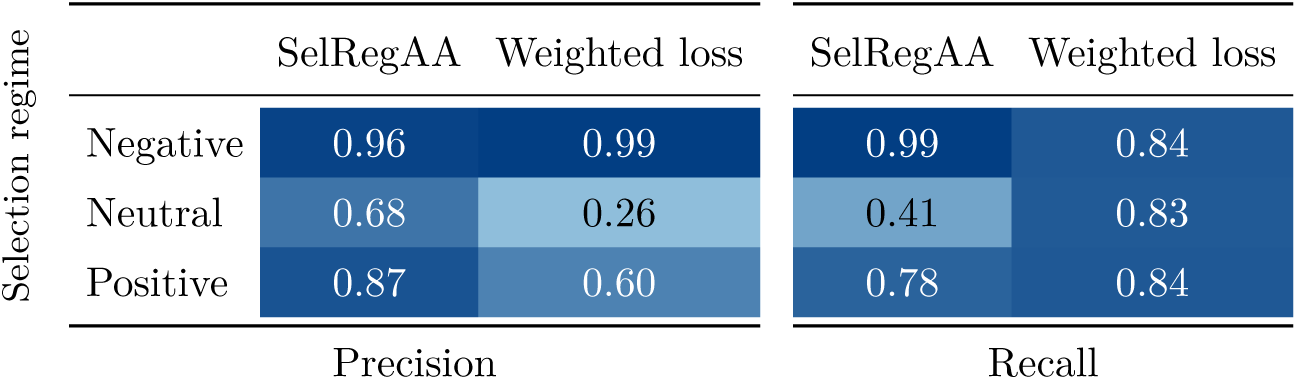
Precision and recall values for SelRegAA (left), and for a model trained with weighted loss (right). The model trained with weighted loss has better recall in the minor categories, and a higher precision in the most frequent category.

We also trained a network with uniform proportions of the three selection regimes, and found that it performed better than SelRegAA on data simulated with uniform proportions, but less well than SelRegAA on data simulated as in SelRegAA’s training data (See Supplementary section 8). Overall, the proportions of selection regimes used for training have a clear effect on the network’s ability to predict selection regimes.

#### 3.2.3 Sensitivity to the trees used

The trees in our training and testing datasets have been simulated as in Nesterenko et al. (2025). Briefly, all of these trees have 50 leaves, and have been scaled to match the distribution of tree diameters observed in the HOGENOM database (Penel et al., 2009). We investigated the sensitivity of SelRegAA to different tree topologies. We obtained 4 empirical phylogenies used in Duchemin et al. (2023). The besnard2009 tree is a 79 leaf phylogeny of the Cyperaceae family of flowering plants, online_rodent is a 32 leaf phylogeny of rodent species, orthomam is a 116 leaf phylogeny of mammalian species, and rubisco is a 179 leaf phylogeny of the Amaranthaceae family of flowering plants. The plant phylogenies have been reconstructed from single gene alignments, while the other two phylogenies have been reconstructed from concatenates. Table 5 presents the *F*_1_ scores for a variety of experiments: we ran inference on the empirical trees (keeping their original branch lengths), on the same trees after rescaling branch lengths to match the branch length of trees used during training, and actual simulated trees with the same number of leaves than the empirical trees. Performance on alignments simulated along the empirical trees (column *Empirical tree*) is systematically lower than the performance obtained on validation data during training (*F*_1_ score on test data of 0.769). Within the empirical trees, we observe that performance is better when the number of sequences is larger, which is consistent with our results obtained above (Table 3). Rescaling the empirical trees to match the tree diameters used in our training simulations improves the performance of the network (column *Scaled empirical tree*). Performance on alignments simulated on new trees with the same number of leaves as the empirical phylogenies, but keeping the scaling as in the training data, results in similar performance (column *Simulated tree*). These results show that both the number of sequences and the scale of the tree have large impacts on the performance of the network, but the topology of the trees, whether they are simulated or empirical, does not seem to matter.

**Table 5:** Performance (*F*_1_ score) of the network on different phylogenies of different lengths. The *F*_1_ score obtained on test data during training is 0.769. When empirical trees are scaled to match the scale of training data, performance improves (column *Scaled empirical tree*). Similarly, performance on new simulated phylogenies with numbers of tips that match the empirical trees, but scale that matches the training data, is also improved (column *Simulated tree*).

| Tree | Empirical tree | Scaled empirical tree | Simulated tree |
| --- | --- | --- | --- |
| besnard2009 | 0.711 | 0.796 | 0.793 |
| online_rodent | 0.624 | 0.712 | 0.729 |
| orthomam | 0.615 | 0.804 | 0.807 |
| rubisco | 0.586 | 0.834 | 0.818 |

**Table 6:** Precision and recall values for our basemodel, CODEML model M2 Bayes Empirical Bayes predictions, CODEML model M8 Bayes Empirical Bayes predictions, and the Persistent Positive Selection test on 10 alignments simulated as during training.

|  |  |  |  |  |  |
| --- | --- | --- | --- | --- | --- |
| Selection regime |  | SelRegAA | CODEML M2 | CODEML M8 | PPS test |
|  | Negative | 0.97 | 0.99 | NA | NA |
|  | Neutral | 0.63 | 0.12 | NA | NA |
|  | Positive | 0.89 | 0.37 | 0.41 | 0.07 |
| Precision |  |  |  |  |  |
| Selection regime |  | SelRegAA | CODEML M2 | CODEML M8 | PPS test |
|  | Negative | 0.98 | 0.62 | NA | NA |
|  | Neutral | 0.43 | 0.79 | NA | NA |
|  | Positive | 0.88 | 0.78 | 0.78 | 0.92 |
| Recall |  |  |  |  |  |

### 3.3 Comparison to standard likelihood-based methods

#### 3.3.1 Performance on data similar to the training data

We compared our network to swMutSel, a method to detect positively selected sites based on the Mutation-Selection framework (Tamuri and dos Reis, 2021), which therefore perfectly fits the simulated data. we additionally tested two *d_N_ /d_S_* methods, the standard method CODEML (Yang, 2007), and a faster implementation, Godon (Davydov et al., 2017). The two *d_N_ /d_S_* methods are at a disadvantage here, because their model does not fit the model that was used to simulate the data. CODEML and Godon implement the same models, and in our tests obtained the same results. We therefore only show results for CODEML, but discuss run times for both implementations. The persistent positive selection test of Tamuri and dos Reis (Tamuri and dos Reis, 2021) fits the simulation model, and therefore provides an interesting comparison to our network. Tests were run on 10 newly simulated alignments, that the base network had not seen before, but simulated with the same pipeline as the training data. CODEML was used on codon sequences to predict whether a site is under negative selection, neutral evolution or positive selection with model M2a (Nielsen and Yang, 1998), and was used to predict positive selection only using the M8 model (Yang et al., 2000). In both cases, prediction relied on the Bayes Empirical Bayes method (Yang et al., 2005). Similarly, the persistent positive selection (PPS) test (Tamuri and dos Reis, 2021) was also applied to predict sites under positive selection only. Overall, SelRegAA has better results than CODEML’s two models (Table 6 ; Supp. Fig. 2), and than the PPS test. In particular, CODEML’s precision for the Neutral evolution and Positive selection classes is much lower than SelRegAA’s precision. CODEML’s recall however is better than SelRegAA’s for the Neutral evolution class, but lower for the two other classes. Overall, for a given recall rate, SelRegAA often offers a better precision than CODEML (Supp. Fig. 2). Prediction of positive selection with CODEML’s model M8 is slightly better than with model M2, but still dominated by SelRegAA. The PPS test for predicting sites under positive selection has very good recall, but very bad precision. This might be linked to optimization difficulties. Indeed, out of the 2500 sites (10 alignments of 250 sites each), the model did not converge for 55 sites, indicating that computation of the PPS test may be hampered by numerical instabilities. Performance metrics were computed on the remaining 2445 sites.

**Figure 2:**
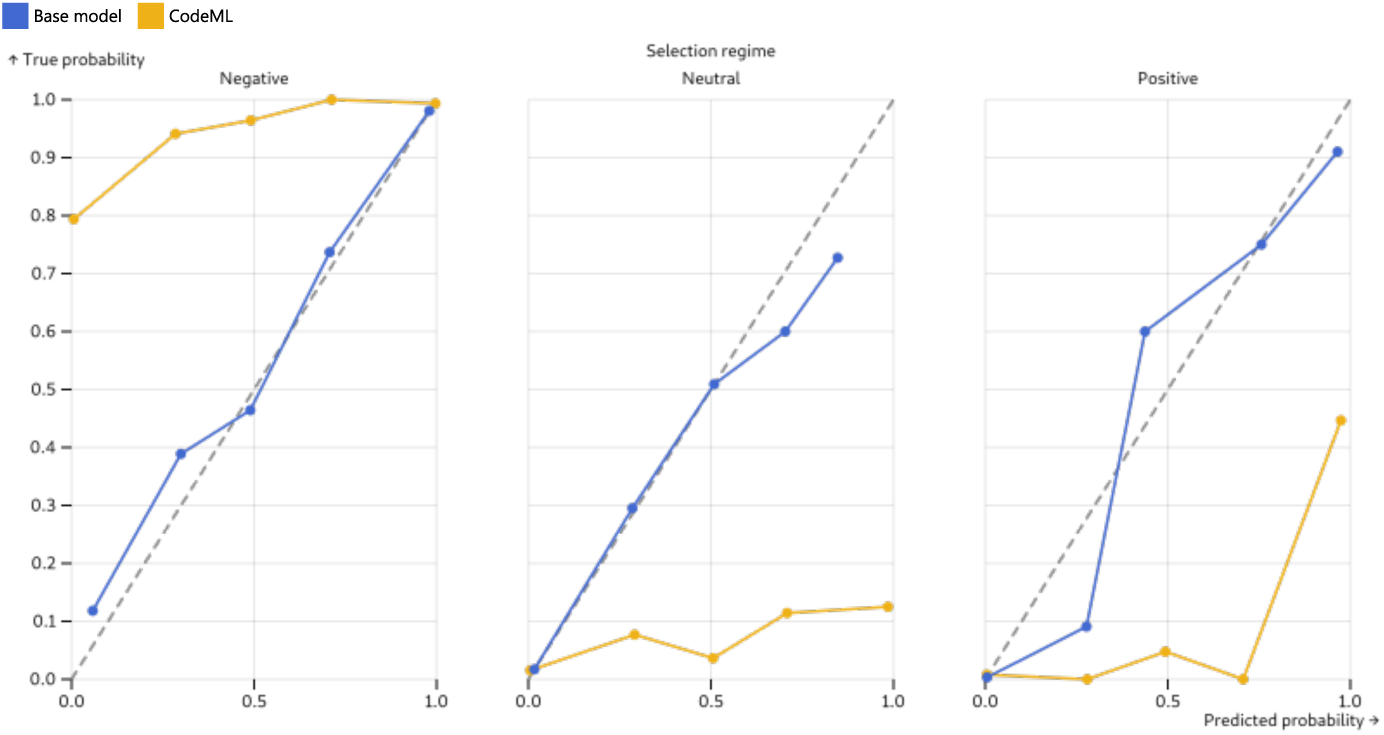
Calibration of SelRegAA and of CODEML. On the x axis, predicted probability that a site is under negative selection (left panel), neutral evolution (middle), or positive selection (right), is compared to the proportion with which a site was indeed in the corresponding selection regime (y axis). A calibrated method is expected to generate points along the y=x regression line.

##### Comparison of computing times

We compared the inference times of the base network, the PPS test, CODEML and Godon, a faster implementation of the codon models implemented in CODEML. Godon was used to perform inference with model M8. Prediction of selection regimes for the 10 alignments required about 3h on 10 CPUs with CODEML, and 20 minutes on 10 CPUs with Godon. This difference is consistent with the range of speed ups reported in Davydov et al. (2017). Computation with the PPS test involves site-wise computations, including 100 replicate computations per individual site to assess significativity. We estimate that running the PPS test for all 10 alignments took the equivalent of more than 700 days of computation, on 10 CPUs. For SelRegAA, inference for all 10 alignments took 5 seconds on a single CPU. The different computations were run on different computers on a cluster; however, the differences in computing time are such that there is no doubt that inference with SelRegAA is orders of magnitude faster than inference with Godon, itself about 10 times faster than CODEML, itself orders of magnitude faster than the PPS test.

Given the better results and runtimes of CODEML over swMutSel, in the following we decided to compare SelRegAA to CODEML rather than swMutSel.

#### 3.3.2 Calibration

We investigated whether the probabilities output by SelRegAA and by CODEML model 0 are calibrated. A probability *p* of an event is calibrated if the event occurs a fraction *p* of the time. Our network outputs for each site a probability that it is under negative, neutral, or positive selection. CODEML uses Bayes Empirical Bayes estimates to compute probabilities for each site to be in one of the three selective regimes (Yang et al., 2005). We used the results on the 10 alignments to compute the frequency with which a prediction is correct and compared it to the probabilities output by the two methods.

Fig. 2 shows that the network’s output probabilities are well calibrated on data similar to its training dataset. Although the neutral and positive categories are much less frequent in the training data than the negative selection category, all three categories are predicted with good calibration. In contrast, CODEML’s M2a Bayes Empirical Bayes probabilities are not well calibrated in all three categories. CODEML is overconfident for neutral and positive categories and, in reverse, is conservative for the negative selection category. The poor calibration of CODEML models M2a is probably because it is measured on MutationSelection simulations that do not correspond to model M2a.

#### 3.3.3 Performance on simulations using different amino acid profiles

Our method was trained and tested on simulations using a limited set of 263 experimentally determined amino acid fitness profiles. We tested its performance on alignments simulated from independently-obtained amino acid profiles and compared it to CODEML. We retrieved amino acid profiles estimated in Latrille et al. (2024) and randomly picked two samples of 250000 profiles (profiles0 and profiles1) from 1000 alignments.

Table 7 shows that the performance of the network suffers markedly when tested on alignments simulated with a different set of profiles, with an *F*_1_ score dropping from 0.769 to about 0.575. In particular, the two classes with the smallest numbers of sites are much less well detected, both in terms of precision and recall.

**Table 7:**
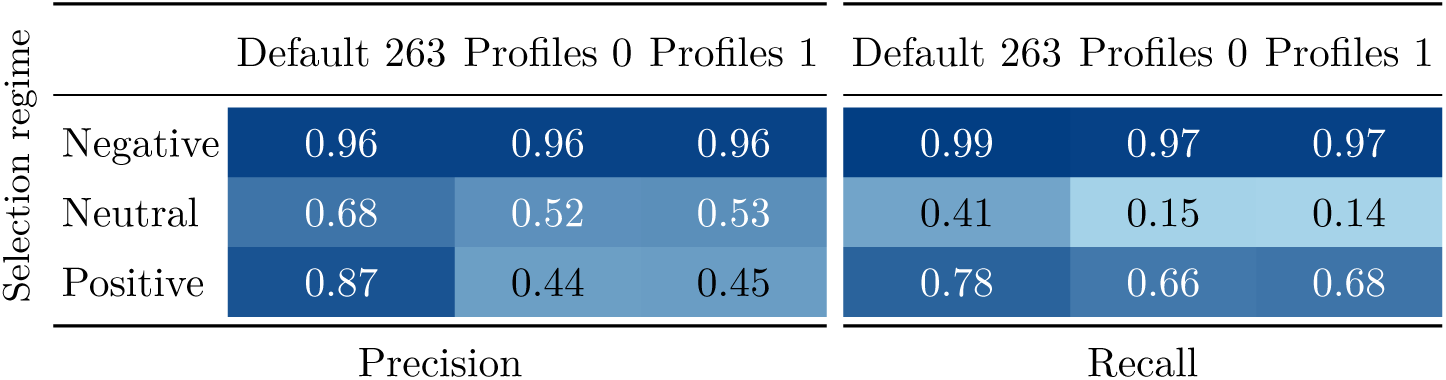
Performance of SelRegAA is markedly decreased on alignments simulated with a different set of amino acid profiles.

A network trained on alignments simulated with profiles0 has reduced performance on alignments simulated with the 263 experimentally determined amino acid fitness profiles (*F*_1_ score of 0.676) (Supp. Table 9). However, performance does not vary whether test alignments have been simulated with profiles0 or profiles1, which both were inferred from large numbers of sites in protein alignments from mammals.

On 10 alignments simulated with profiles0, comparison between CODEML and SelRegAA trained on the 263 default profiles shows that the performance gap between the two models has decreased with *F*_1_ scores of 0.539 and 0.548 respectively (compared to 0.492 and 0.791 on data simulated with the 263 default profiles), although SelRegAA keeps a small edge. The two methods, however, exhibit different behaviours (Table 8). For example, they have opposite positions on the precision-recall trade-off for the neutral category, with SelRegAA showing a bad recall but better precision, while CODEML shows a bad precision with better recall.

**Table 8:**
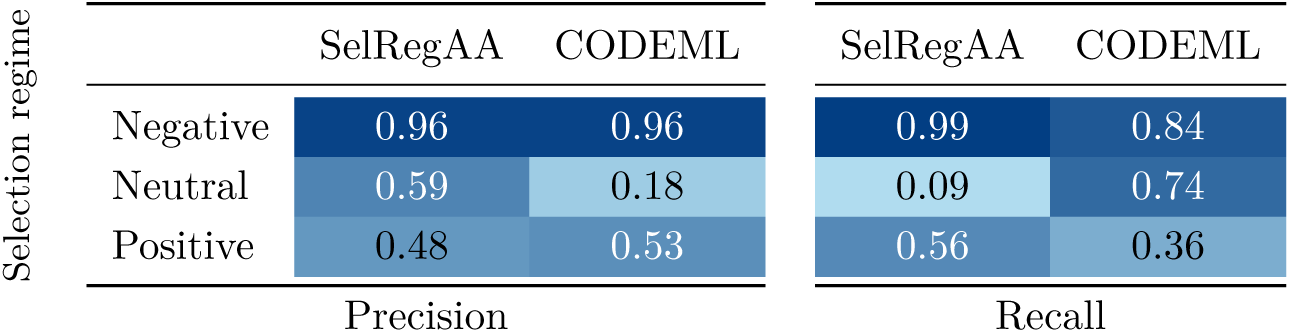
Comparison between CODEML and SelRegAA (trained on the default 263 profiles) on alignments simulated with profiles0.

**Table 9:**
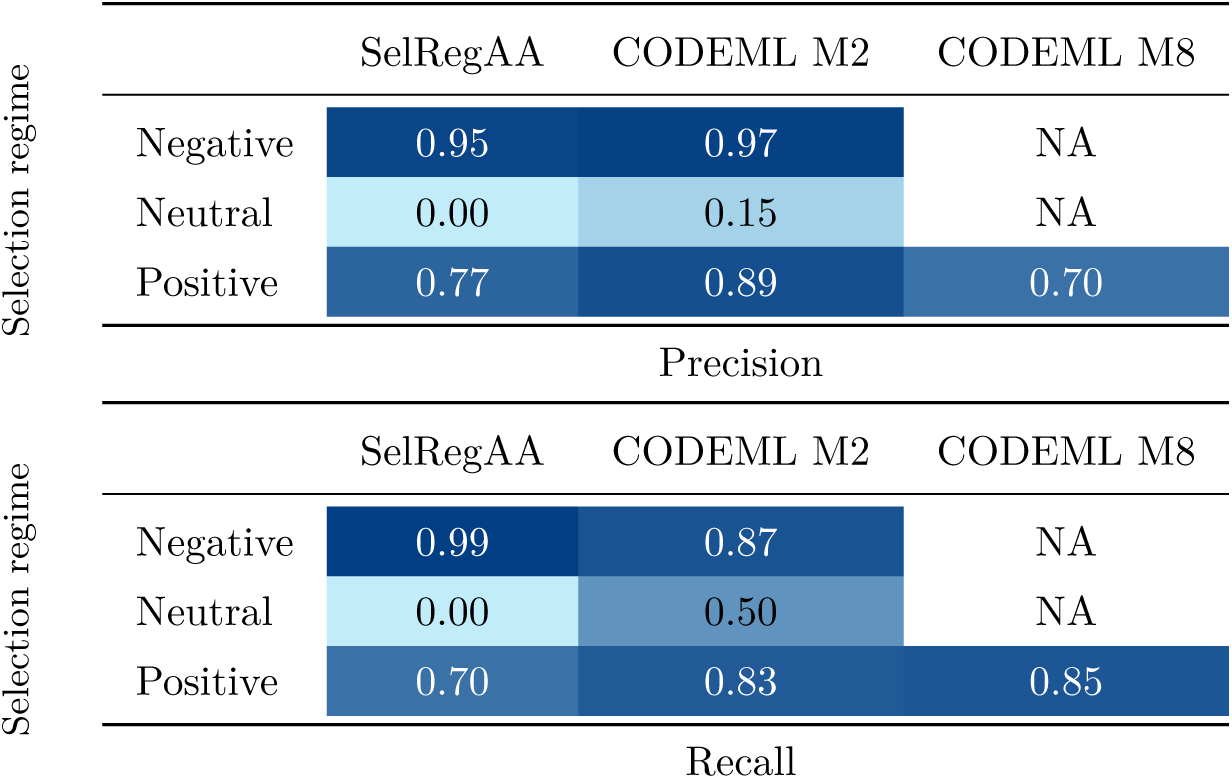
Precision and recall values for our basemodel, CODEML model M2 Bayes Empirical Bayes predictions and CODEML model M8 Bayes Empirical Bayes predictions on 10 alignments of 250 codons simulated under a *d*_N_ */d*_S_ model.

This drop in performance for SelRegAA is also seen in the calibration of the probabilities (Fig. 3). In particular, SelRegAA is now overconfident in its  prediction of sites in the positive selection category, with a behaviour that is now similar to CODEML.

**Figure 3:**
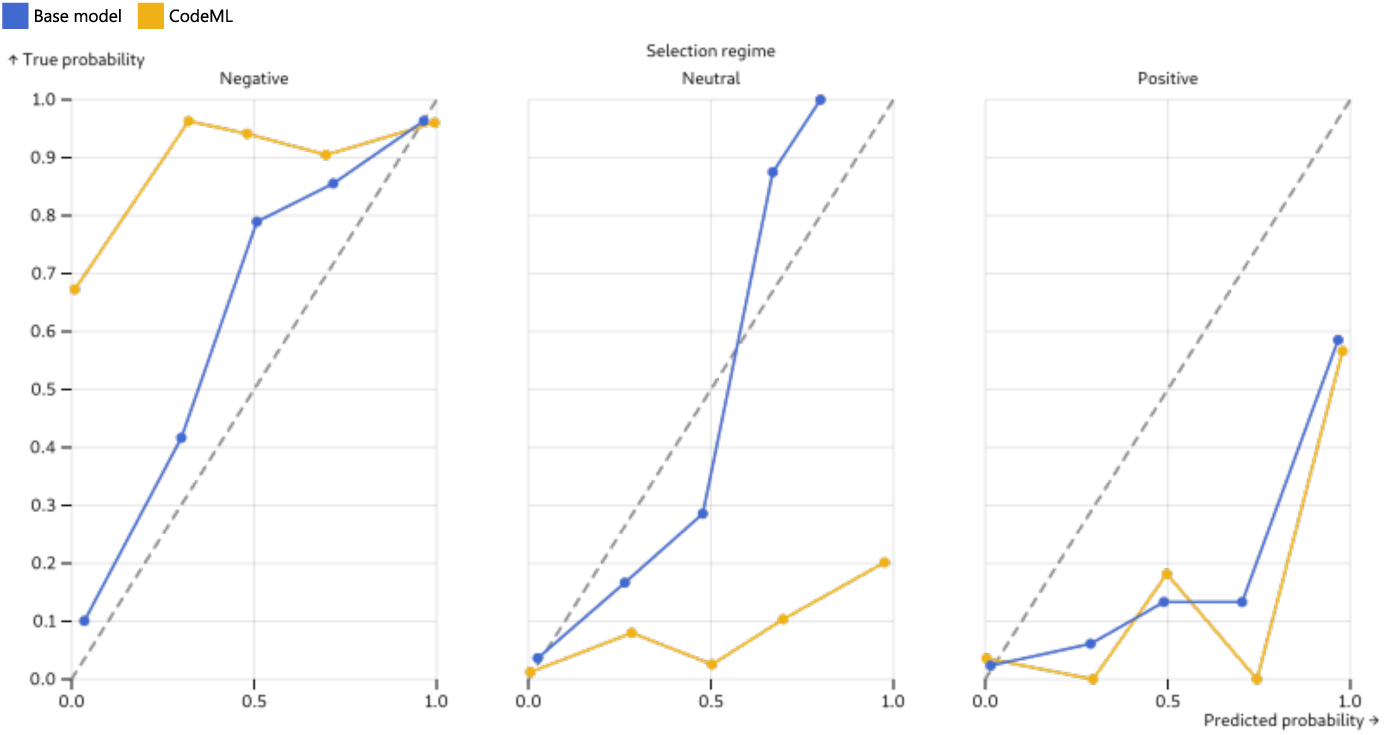
Calibration of CODEML and the base model on simulations using amino acid profiles different from the training data

#### 3.3.4 Performance on simulations using a *d_N_ /d_S_* model

On alignments simulated under a *d_N_ /d_S_* model, SelRegAA remained accurate for sites under negative selection, with a precision of 0.95, a recall of 0.99 and an *F*_1_ score of 0.97 (Table 9). Its performance for positive selection was also encouraging, with a precision of 0.77, a recall of 0.70 and an *F*_1_ score of 0.73, showing that the network still performed well even though the data were generated under a model not used during training. SelRegAA did not, however, correctly identify neutral sites (*F*_1_ score of 0). CODEML M2 Bayes Empirical Bayes inference performed better for both neutral and positively selected sites (*F*_1_ scores of 0.23 and 0.86, respectively). CODEML M8 Bayes Empirical Bayes inference achieved the highest recall for positive selection, but had the lowest precision, resulting in a *F*_1_ score of 0.76. Because SelRegAA was trained exclusively on Mutation-Selection simulations, its decreased performance is likely due to the difference between the training and test models. Calibration of the predicted probabilities on these *d_N_ /d_S_*simulations varies across selection regimes and methods (Fig. 4). SelRegAA’s predicted probability for neutral evolution remains close to zero even when a site is truly neutral, and it is overconfident when predicting negative selection. CODEML remains overconfident for neutral evolution and conservative for negative selection, as already observed on Mutation-Selection simulations (Fig. 2). For positive selection, CODEML’s predictions remain well calibrated, whereas SelRegAA is conservative when its predicted probability is low but becomes overconfident when it is high.

**Figure 4:**
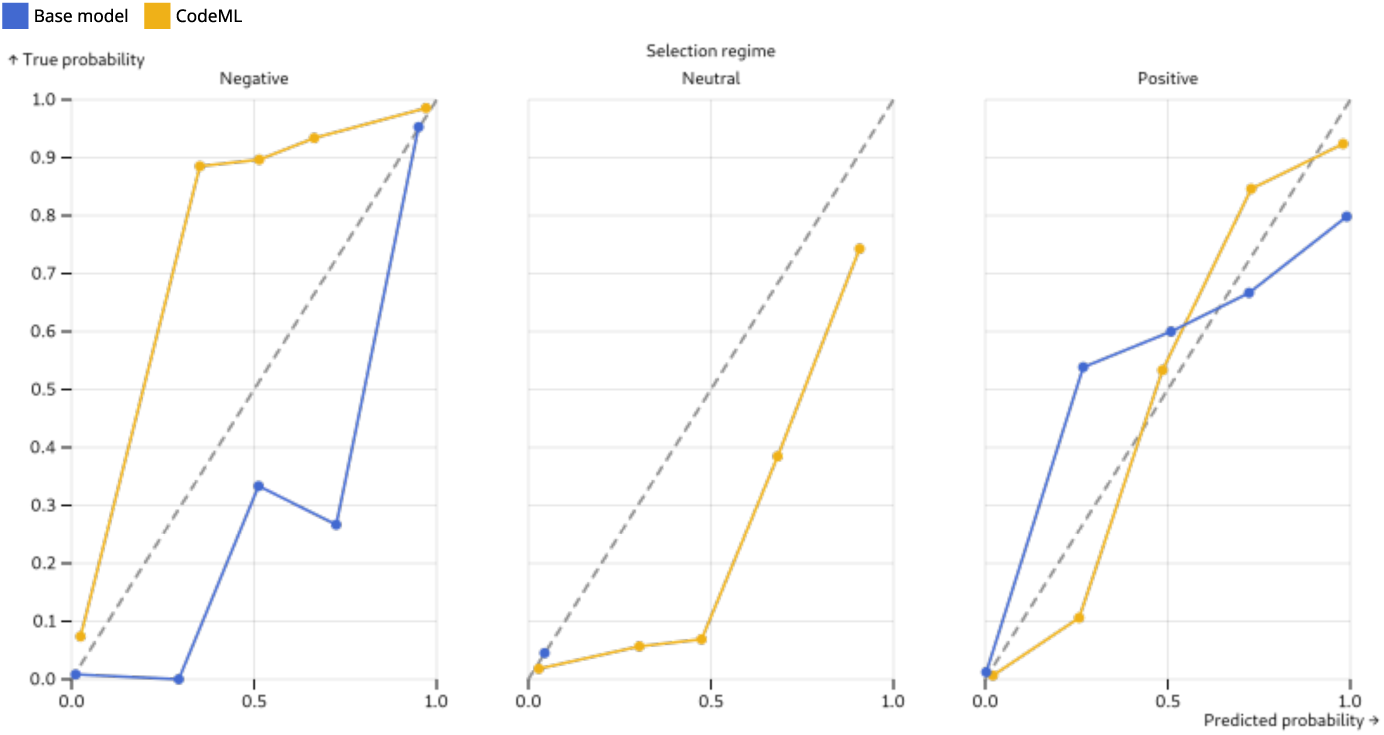
Calibration of CODEML and the basemodel on simulations using a *d*_N_ */d*_S_ model.

#### 3.3.5 Comparison of methods using empirical data

Inference of positive selection with SelRegAA, CODEML M8 Bayes Empirical Bayes inference, and the *ω*-based proxy derived from the PPS model (Fig 5) was generally congruent on empirical data. However, SelRegAA tended to detect more sites under positive selection than the two other methods. This could be consistent with the higher recall of SelRegAA for positive sites in simulations.

**Figure 5:**
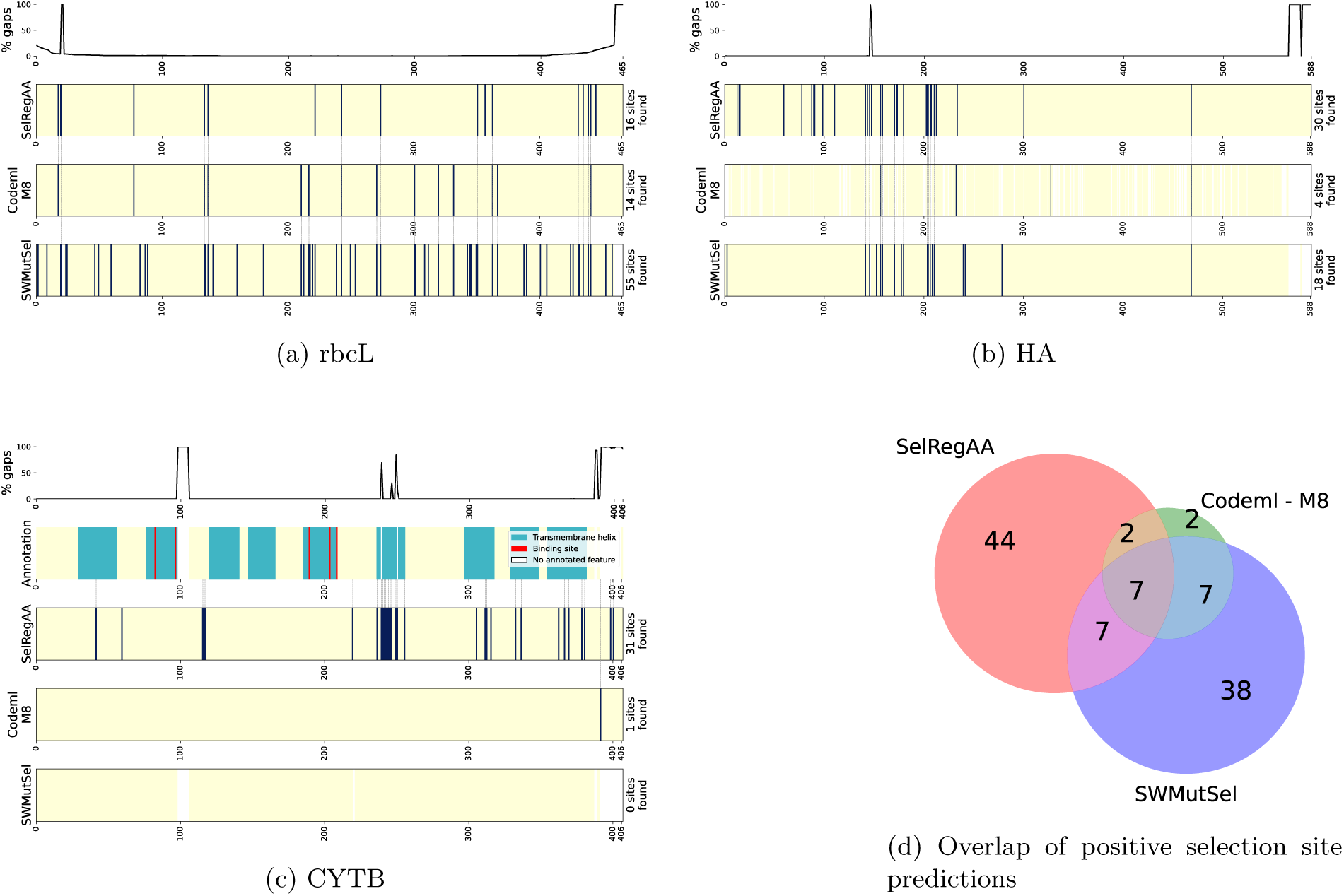
Site-wise comparison of positive selection predictions for the empirical rbcL, HA and CYTB alignments. (a-c) The upper track shows the percentage of gaps at each alignment position. The following tracks show sites predicted to be under positive selection by SelRegAA, CODEML M8 Bayes Empirical Bayes inference, and the *ω*-proxy of the PPS test obtained with *λ*= 0.001. For CYTB alignment (c) an additional track shows the annotation of each position, retrieved from Uniprot, accession Q6VHH3 (The UniProt Consortium, 2025). (d) Overlap of the sites predicted to be under positive selection by the three methods, pooled across the three alignments. Only sites analysed by all three methods are included.

For the rbcL alignment, 14 of the 16 sites predicted to be under positive selection by SelRegAA were also detected by CODEML M8 or by the PPS-test proxy. Most SelRegAA predictions for this alignment were independently supported by at least one other approach. SelRegAA also recovered 5 of the 12 sites jointly identified by CODEML and the PPS-test proxy. For the HA alignment, 11 of the 30 SelRegAA predictions were also detected by at least one of the other methods. SelregAA also recovered both sites jointly identified by CODEML and the PPS-test proxy.

The results differed for the CYTB alignment, for which none of the 31 sites predicted by SelRegAA was supported by other methods. In this alignment, the PPS-test proxy detected no site under positive selection and CODEML M8 Bayes Empirical Bayes inference detected only 1 site under positive selection. Notably, the sites predicted by SelRegAA were more often located within the transmembrane helices annotated in UniProt (The UniProt Consortium, 2025) than in the rest of the protein.

SelRegAA provided predictions for all sites, whereas CODEML ignored sites with gaps in the HA alignment, and Tamuri and dos Reis (2021) did not provide swMutSel predictions for all sites. To compare the three methods across all alignments, we pooled their predictions and restricted the comparison to sites analysed by all three methods (Fig 5d). For each pair of methods, we calculated the proportion of sites detected by both methods among all sites detected by at least one of the two. This proportion was 13% between SelRegAA and the PPS-test proxy, 13% between SelRegAA and CODEML M8, and 22% between CODEML M8 and the PPS-test proxy. CODEML M8 and the PPS-test proxy therefore agreed more with each other than they did with SelRegAA.

Overall, the empirical results shows that SelRegAA predicts more sites under positive selection than the two conventional approaches, but that several sites that it predicts are also found by at least one of the two other methods.

## Discussion

In this manuscript, we have presented our work to develop and train a neural network to predict selective regimes at amino acid alignment sites. We have characterized how the performance of the network varied with features of the network architecture and with characteristics of the training data. These results can be used as guidance for training these types of neural networks for site-wise inference on multiple sequence alignments. In particular, we found that varying the embedding size, the number of attention blocks or attention heads had little effect on performance. However, handling pairs of sequences and using larger training datasets improved our results. These two approaches however have shortcomings: training on a larger dataset entails a larger computational footprint, and the model working with pairs of sequences both requires longer training and has a much larger memory footprint than the model working with sequences. As a result, we focused our evaluations on a version of our network that was not the top performer, but that offered a good trade-off between computational footprint and accuracy.

The results obtained by SelRegAA on simulated data are very promising. Comparison with swMutSel and CODEML showed that the network achieved better performance, at a fraction of the computing time, when the model used to generate the test data matches that used to generate the training data. SelregAA’s good performance is surprising because, contrary to its competitors, it does not have access to the phylogeny of the sequences, and to synonymous changes, which provide information on the neutral rate of evolution. This suggests that a similar network, trained on codon sequences, could achieve even higher performance. The fact that swMutSel performs less well than CODEML suggests that swMutSel may get trapped in local optima during parameter optimization. In the comparison with CODEML, SelregAA’s better performance is probably the result of (i) a mismatch between the model used to simulate test data and the model underlying CODEML and (ii) the fact that the network performs Bayesian inference and so benefits from the information contained in the prior. Regarding (i), our simulations rely on the mutation-selection model, not on a *d_N_ /d_S_* model. This causes a decreased performance of CODEML compared to its performance on data simulated in the *d_N_ /d_S_* framework, as can be seen from the comparison of Tables 6 and 9. Similarly, the probabilities output by the Bayes Empirical Bayes estimates provided by CODEML (Yang et al., 2005) were less well calibrated on data simulated in the mutation-selection framework than on data simulated in the *d_N_ /d_S_* framework (compare Figs. 2 and 4). However, the mutation-selection model appears to be more realistic than *d_N_ /d_S_* models (Jones et al., 2019; Rodrigue et al., 2020), and past work on protein sequences shows that taking into account amino acid composition heterogeneity between sites improves model adequacy (Lartillot et al., 2007). This literature implies that it could be important to use methods that perform well on data simulated in the Mutation-Selection framework when analyzing empirical data. Regarding (ii): training the network on simulated data amounts to teaching it to perform Bayesian inference (Nesterenko et al., 2025). For this reason, when the neural network operates with a correctly specified prior, it has an advantage over CODEML (Table 6), with a better calibration (Fig. 2). Conversely, when it is used on data simulated in the *d_N_ /d_S_* framework, it is at a disadvantage, as can be seen Table 9, with a worse calibration (Fig. 4). In between these two extremes, Table 7 shows that using amino acid equilibrium profiles that differ from the training data results in decreased performance, and decreased calibration 3, for an overall performance that is similar or slightly better than CODEML.

Our results obtained with different amino acid profiles, with different phylogenies, or with different proportions of selection regimes, show that the performance of the model is very dependent on the match between training and testing data. Once again, we suspect this is because the training of a neural network on simulations amounts to building a model that performs Bayesian inference (Nesterenko et al., 2025). The resulting network therefore encapsulates a prior distribution over all parameters of the simulation model, which can result in altered performance if there are mismatches between the model that generated the training data and the model that generated the test data. This means that when analyzing empirical data, much attention must be devoted to generating realistic data for the training. Here we have anchored the tree diameters on an empirical tree distribution, and the amino acid profiles on empirical data. Unfortunately however, it is certain that the empirical distributions that we have used do not cover the whole range of parameter values that would be needed to fit different types of alignments. Alignments from different groups of species will differ in their size, their evolutionary depth, their GC content, or the selective pressures that have applied on them. Training a network that could perform equally well on any type of biological alignment is an important challenge. Perhaps a feasible approach would be to train several networks to cover different areas of the parameter space, and develop a method to choose the most relevant network depending on characteristics of the alignment under study. Another solution would be to train a network *de novo* for a given dataset. Our results show that the training of our network on a GPU plus the inference on 10 alignments on a single CPU took about the same amount of time as running CODEML on these 10 alignments on 10 CPUs. When inference needs to be performed at the genomic scale, for thousands of gene alignments, training de novo, on simulations that resemble the empirical data, provides a lower footprint than using CODEML. Our open source code can be deployed for training or fine-tuning a model on new simulations. However, choosing parameter values that generate datasets that are similar to empirical data remains an area of research (see Trost et al. (2024) and, for gap patterns, Ashkenazy et al. (2017)). For the time being, analysis of 3 empirical alignments showed some overlap between SelRegAA and CODEML and swMutSel in their identification of positively selected sites, but also showed that SelRegAA tended to detect a larger number of positive sites (Fig. 5). This could be a result of SelRegAA’s tendency to be over-confident in calling positive selection in simulations that don’t match its training data, as was seen in the *d_N_ /d_S_* simulations Fig. 4.

This work further suggests that similar neural networks could be used to predict other features of amino acid sites. For instance, a network could be used to predict sites undergoing shifts in the direction of selection associated to phenotype changes (Duchemin et al., 2023). To do so, the network should take as additional input phenotypic states at each leaf of the tree, or a phylogenetic tree annotated with phenotypic states at each node. Beyond site-wise prediction, similar networks could also predict model parameters that apply to an entire alignment, such as a matrix of nucleotide substitutions, or an average *d_N_ /d_S_*. Because the transformer architecture at the heart of our model can capture dependencies between sites of the alignment, one could also design a model to learn parameters of models that capture co-evolution between sites, such as the Direct Coupling Analysis model (Morcos et al., 2011).

## Conclusion

In this work, we have shown that a neural network trained on simulated amino acid sequences could predict selection pressures acting on individual protein sites. It performed better than state-of-the-art methods across datasets that resemble its training data, but can perform less well when there is a difference between training and test data. Looking ahead, this shows that a similar network working on codon sequences, and trained on data that resembles empirical data, could markedly outperform maximum likelihood approaches in speed and accuracy to characterize selective pressures acting on sites. Further, it paves the way for the inference of other aspects of molecular evolution.

## Acknowledgements

This work was funded by the Agence Nationale de la Recherche (PIECES ANR20-CE45-0017, and Deelogeny ANR-23-CE45-0027). This work was performed using the computing facilities of the CC LBBE/PRABI and of the CC IN2P3.

## Notes

https://gitlab.in2p3.fr/deelogeny/wp4/selregaa

### Competing Interest Statement

The authors have declared no competing interest.

### Summary of Updates

This version has been edited to address a second round of reviews by independent researchers. It includes new experiments on simulated data in the dN/dS framework, and on experimental alignments, and text changes.

