## Supplementary information for "Characterization of selective pressures acting on protein sites with Deep Learning"

### Supplementary Material for Characterization of selective pressures acting on protein sites with Deep Learning

#### 1 Sequence to embedding transformation

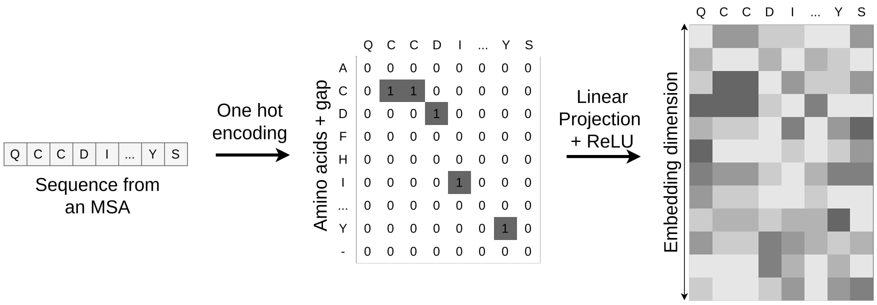


Supplementary Figure 1: A sequence extracted from a multiple sequence alignment (MSA) is first encoded using a one-hot representation, where each amino-acid is represented by a vector of size 21, corresponding to the 20 amino acids plus one gap symbol. This representation is then projected into an embedding space of dimension 32 through a linear transformation followed by a non-linear activation (ReLU).

#### 2 Replicability

| Selection regime | Base model | Base model 2 | Base model 3 | Base model 4 |
|---|---|---|---|---|
| Negative | 0.96 | 0.96 | 0.96 | 0.96 |
| Neutral | 0.68 | 0.66 | 0.67 | 0.67 |
| Positive | 0.87 | 0.87 | 0.88 | 0.86 |

Precision

| Selection regime | Base model | Base model 2 | Base model 3 | Base model 4 |
|---|---|---|---|---|
| Negative | 0.99 | 0.99 | 0.99 | 0.99 |
| Neutral | 0.41 | 0.40 | 0.40 | 0.40 |
| Positive | 0.78 | 0.75 | 0.76 | 0.77 |

Recall

Supplementary Table 1: Performance across training replicates of the base network architecture. Statistics shown were computed on the validation data used during training.

#### 3 Effect of training on a larger training set

| Selection regime | Precision: 50 000 alignments | Precision: Base model | Recall: 50 000 alignments | Recall: Base model |
|---|---|---|---|---|
| Negative | 0.96 | 0.96 | 0.99 | 0.99 |
| Neutral | 0.69 | 0.68 | 0.42 | 0.41 |
| Positive | 0.92 | 0.87 | 0.82 | 0.78 |

Supplementary Table 2: Precision and recall for base model (right) and for the network trained on a larger data set of size 50000 alignments (left).

#### 4 Effect of the embedding size

| Selection regime | Base model | Embed dim 16 | Embed dim 48 | Base model | Embed dim 16 | Embed dim 48 |
|---|---|---|---|---|---|---|
| Negative | 0.96 | 0.96 | 0.96 | 0.99 | 0.99 | 0.99 |
| Neutral | 0.68 | 0.66 | 0.68 | 0.41 | 0.38 | 0.41 |
| Positive | 0.87 | 0.85 | 0.89 | 0.78 | 0.74 | 0.77 |
| | Precision | | | Recall | | |

Supplementary Table 3: Effect of the embedding size on the performance of the network. Left: base model. Middle: the embedding size has been shrunk to 16 instead of 32. Right: the embedding size has been increased to 48.

#### 5 Effect of the number of attention blocks

| Selection regime | 2 blocks | 4 blocks | 8 blocks | Base model |
|---|---|---|---|---|
| Negative | 0.96 | 0.96 | 0.96 | 0.96 |
| Neutral | 0.66 | 0.68 | 0.67 | 0.68 |
| Positive | 0.85 | 0.87 | 0.87 | 0.87 |

Precision

| Selection regime | 2 blocks | 4 blocks | 8 blocks | Base model |
|---|---|---|---|---|
| Negative | 0.99 | 0.99 | 0.99 | 0.99 |
| Neutral | 0.37 | 0.39 | 0.39 | 0.41 |
| Positive | 0.73 | 0.76 | 0.76 | 0.78 |

Recall

Supplementary Table 4: Effect of the number of attention blocks on the performance of the network. Left: model with 2 attention blocks. Second left: model with 4 attention blocks. Third left: model with 8 attention blocks. Fourth: base model.

#### 6 Effect of the number of attention heads

| Selection regime | 2 heads | 6 heads | Base model | 2 heads | 6 heads | Base model |
|---|---|---|---|---|---|---|
| Negative | 0.96 | 0.96 | 0.96 | 0.99 | 0.99 | 0.99 |
| Neutral | 0.65 | 0.66 | 0.68 | 0.39 | 0.42 | 0.41 |
| Positive | 0.85 | 0.87 | 0.87 | 0.74 | 0.77 | 0.78 |
| | Precision | | | Recall | | |

Supplementary Table 5: Effect of the number of attention heads on the performance of the network. Left: model with 2 attention heads. Middle: model with 6 attention heads. Right: base model.

#### 7 Effect of the number of sites

| Selection regime | 400 sites | Base model | 400 sites | Base model |
|---|---|---|---|---|
| Negative | 0.96 | 0.96 | 0.99 | 0.99 |
| Neutral | 0.68 | 0.68 | 0.38 | 0.41 |
| Positive | 0.85 | 0.87 | 0.76 | 0.78 |
| | Precision | | Recall | |

Supplementary Table 6: The performance of the trained network does not improve as it is tested in alignments with more sites.

#### 8 Effect of the proportions of sites in each selective regime

We trained our network on 10000 alignments simulated with uniform proportions of negative selection/neutral evolution/positive selection sites ([0.33;0.33;0.33]). Performance on data simulated with the same uniform proportions is improved for this model compared to SelRegAA:

| Selection regime | Precision: SelRegAA | Precision: Uniform | Recall: SelRegAA | Recall: Uniform |
|---|---|---|---|---|
| Negative | 0.45 | 0.84 | 0.99 | 0.83 |
| Neutral | 0.90 | 0.82 | 0.26 | 0.87 |
| Positive | 0.97 | 0.93 | 0.50 | 0.89 |

Supplementary Table 7: A network trained with uniform proportions of selective regimes performs better than SelRegAA on data simulated the same way.

Conversely, this network trained on data with uniform proportions of selective regimes performs less well than SelRegAA when data is simulated with selection regime proportions that match SelRegAA's training data.

| Selection regime | Precision: SelRegAA | Precision: Uniform | Recall: SelRegAA | Recall: Uniform |
|---|---|---|---|---|
| Negative | 0.96 | 0.99 | 0.99 | 0.55 |
| Neutral | 0.68 | 0.08 | 0.41 | 0.43 |
| Positive | 0.87 | 0.19 | 0.78 | 0.92 |

Supplementary Table 8: SelRegAA performs better than a network trained with uniform proportions of selective regimes on data simulated as in its training data.

### 9 Precision-recall curves per selection regime

Detailed investigation of the precision-recall curves (Fig. 2) shows that SelRegAA achieves better recall and precision than CodeML.

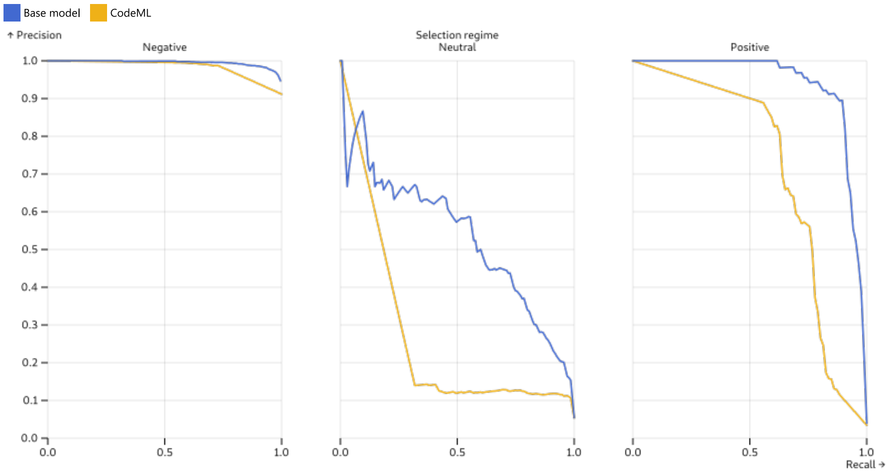


Supplementary Figure 2: precision-Recall curves for the base model (yellow) and for codeml (blue).

#### 10 Performance of the model across profile sets

| Selection regime | Precision | | | Recall | | |
|---|---|---|---|---|---|---|
| | Default 263 | Profiles 0 | Profiles 1 | Default 263 | Profiles 0 | Profiles 1 |
| Negative | 0.96 | 0.96 | 0.96 | 0.96 | 0.99 | 0.99 |
| Neutral | 0.42 | 0.81 | 0.81 | 0.35 | 0.63 | 0.61 |
| Positive | 0.64 | 0.85 | 0.85 | 0.73 | 0.61 | 0.63 |

Supplementary Table 9: Performance of the model trained on $profiles_0$ is markedly decreased on alignments simulated with the 263 default amino acid profiles, but similar on $profiles_0$ and $profiles_1$ alignments.
